# RNA-mediated ribonucleoprotein assembly controls TDP-43 nuclear retention

**DOI:** 10.1101/2023.08.06.552215

**Authors:** Patricia M. dos Passos, Erandika H. Hemamali, Lohany D. Mamede, Lindsey R. Hayes, Yuna M. Ayala

## Abstract

TDP-43 is an essential RNA-binding protein strongly implicated in the pathogenesis of neurodegenerative disorders characterized by cytoplasmic aggregates and loss of nuclear TDP-43. The protein shuttles between nucleus and cytoplasm, yet maintaining predominantly nuclear TDP-43 localization is important for TDP-43 function and for inhibiting cytoplasmic aggregation. We previously demonstrated that specific RNA binding mediates TDP-43 self-assembly and biomolecular condensation, requiring multivalent interactions via N- and C-terminal domains. Here, we show that these complexes play a key role in TDP-43 nuclear retention. TDP-43 forms macromolecular complexes with a wide range of size distribution in cells and we find that defects in RNA binding or inter-domain interactions, including phase separation, impair the assembly of the largest species. Our findings suggest that recruitment into these macromolecular complexes prevents cytoplasmic egress of TDP-43 in a size-dependent manner. Our observations uncover fundamental mechanisms controlling TDP-43 cellular homeostasis, whereby regulation of RNA-mediated self-assembly modulates TDP-43 nucleocytoplasmic distribution. Moreover, these findings highlight pathways that may be implicated in TDP-43 proteinopathies and identify potential therapeutic targets.

## INTRODUCTION

TDP-43 (TAR DNA binding protein), a highly conserved and essential RNA binding protein in multicellular eukaryotes, is central in the pathogenesis of neurodegenerative disorders. Accumulation of TDP-43 aggregates and loss of nuclear protein localization are hallmark features of almost all amyotrophic lateral sclerosis (ALS) and ∼50% of frontotemporal dementia (FTD) patients (Arai *et al*, 2006; Neumann *et al*, 2007; Neumann *et al*, 2006). In addition, TDP-43 pathology is a signature feature of an Alzheimer’s disease (AD)-related dementia affecting the oldest population, termed limbic-predominant age-related TDP-43 encephalopathy (LATE) (Nelson *et al*, 2019). The majority of TDP-43 inclusions are observed in the cytoplasm and often correspond to loss of nuclear localization. Structural defects in the nuclear pore complex linked to ALS and FTD promote TDP-43 cytoplasmic mislocalization, suggesting that aberrant nucleocytoplasmic transport impacts TDP-43 homeostasis in disease (Coyne & Rothstein, 2022). Aberrant accumulation of cytoplasmic TDP-43 is linked to increased protein aggregation and is thought to promote depletion of the nuclear pool of functional TDP-43 in affected cells (Gasset-Rosa *et al*, 2019; Mann *et al*, 2019; Winton *et al*, 2008). Maharana et al. proposed that the relatively lower levels of RNA in the cytoplasm is a major factor contributing to aggregation of TDP-43 and other RNA binding proteins in this compartment (Maharana *et al*, 2018). While in the nucleus, RNA binding may prevent TDP-43 misfolding despite the higher concentration of TDP-43. Data from our group and others support this model by showing that specific RNA binding decreases TDP-43 aggregation and promotes liquid properties during phase separation (Grese *et al*, 2021; Mann *et al*., 2019). Therefore, defects in TDP-43 nuclear retention may strongly contribute to pathogenesis by increasing the cytoplasmic accumulation of the protein. At the same time, loss of nuclear localization decreases TDP-43 nuclear functions, such as control of gene expression (Polymenidou *et al*, 2011; Rot *et al*, 2017; Tollervey *et al*, 2011; Wood *et al*, 2020), as detected in neurons extracted from ALS/FTD patient brain devoid of nuclear TDP-43 (Liu *et al*, 2019). Indeed, dysregulation of TDP-43 targets is observed in patients (Brown *et al*, 2022; Ling *et al*, 2015; Ma *et al*, 2022; Melamed *et al*, 2019), supporting the idea that loss of nuclear localization strongly impairs physiological TDP-43 function contributing to disease.

TDP-43 shuttles continuously between the nuclear and cytoplasmic compartments, but is normally predominantly nuclear as shown by fractionation experiments and cell imaging (Ayala *et al*, 2008). TDP-43 undergoes active nuclear import mediated by a basic bipartite nuclear localization signal (NLS) (**Fig.1A**) and binding to importins α/β. (karyopherins α/β)(Nishimura *et al*, 2010; Winton *et al*., 2008). In addition, alternative mechanisms may also actively or passively promote nuclear entry (Ederle *et al*, 2018; Pinarbasi *et al*, 2018; Timney *et al*, 2016). A nuclear export signal was previously proposed to interact with exportin 1 (XPO/CRM1)(Winton *et al*., 2008); however, multiple lines of evidence indicate that TDP-43 passively exits the nucleus in a size-dependent manner (Archbold *et al*, 2018; Duan *et al*, 2022; Ederle *et al*., 2018; Pinarbasi *et al*., 2018) through nuclear pore channels, which increasingly exclude cargoes above 30-60 kDa in size (Mohr *et al*, 2009; Timney *et al*., 2016). Recent findings suggest that binding to nuclear RNAs restricts TDP-43 availability for passive exit through nuclear pore channels (Duan et al., 2022). Thus, TDP-43 incorporation into high molecular weight nuclear ribonucleoproteins (RNPs) may be a key mechanism promoting TDP-43 nuclear accumulation.

**Figure 1.**
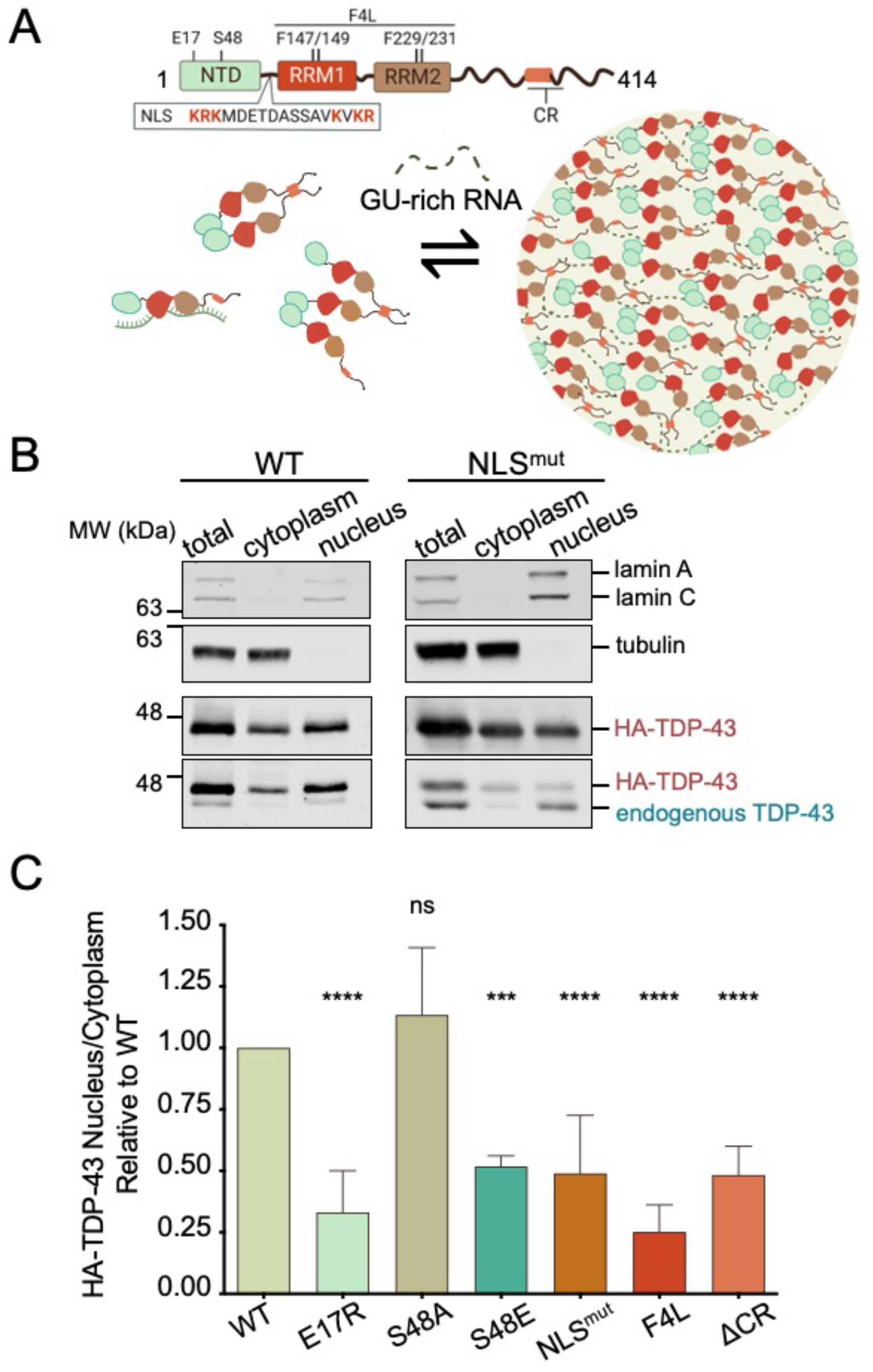
Defects in RNA-mediated TDP-43 self-assembly decrease nuclear localization. A) TDP-43 domain organization and model of RNA-mediated TDP-43 self-assembly involving multivalent interactions through the N-terminal domain (NTD) oligomerization and C-terminal assembly. Mutations in the NTD, RNA recognition motifs (RRMs), and deletion of the conserved α-helical region (ΔCR) in the C-tail (a.a. 316-343) are highlighted. The basic residues (red) in the nuclear localization signal (NLS) were substituted to generate NLS^mut^, used as control. B) Immunoblot of total, cytoplasmic and nuclear fractions of HEK293^HA-TDP-43^ cells. Lamin A/C and tubulin were used as nuclear and cytoplasmic controls, respectively. C) HA-TDP-43 nuclear to cytoplasmic signal ratio (N/C) quantified from immunoblotting, normalized to WT in each independent experiment. Mean ± SD is shown for 6 independent replicates. ns, not significant, ***p<0.001, ****p<0.0001 multiple comparison one-way ANOVA with Tukey’s test.

TDP-43 is a heterogeneous ribonucleoprotein (hnRNP) composed of multiple independently folded domains and a mostly disordered C-terminal region (**Fig 1A**). The N-terminal domain (NTD) resembles ubiquitin-like Dishevelled/Axin (DIX) domains and forms dimers/oligomers through head-to-tail interactions (Afroz *et al*, 2017; Mompean *et al*, 2016). Two tandem RNA recognition motifs (RRMs) bind RNA with strong preference for GU-rich sequences, which are recognized with affinity in the low nanomolar range (Ayala *et al*, 2005; Lukavsky *et al*, 2013). The C-terminal domain (CTD) is a low complexity sequence region also referred to as prion-like domain. The isolated CTD undergoes liquid-liquid phase separation (LLPS) in vitro (Conicella *et al*, 2016) and is believed to largely contribute to TDP-43 condensation in cells (Schmidt & Rohatgi, 2016). In particular, an evolutionarily conserved region (CR, a.a. 320-343) in the CTD that forms a partial α-helical structure (Lim *et al*, 2016) mediates homotypic assembly and LLPS in vitro and in cells (Conicella *et al*., 2016; Schmidt & Rohatgi, 2016). This process of biomolecular condensation is increasingly believed to mediate the formation of protein and RNA-rich assemblies or RNP complexes (Brangwynne *et al*, 2009; Strom & Brangwynne, 2019). In cells, TDP-43 forms and is recruited to RNPs, such as Cajal bodies, and stress granules (Colombrita *et al*, 2009; Tsuiji *et al*, 2013). We recently showed that RNA binding to extended GU-rich sequences promotes TDP-43 biomolecular condensation in vitro and in mammalian cell lysate reconstitution assays (Grese *et al*., 2021). Consistent with the strong contribution of biopolymers and multivalent interactions to the formation of biomolecular condensates (Shin & Brangwynne, 2017), we found that RNA-mediated TDP-43 condensates require NTD and CTD-mediated interactions, as shown in our model (**Fig. 1A**). These TDP-43 assemblies are necessary for the regulation of RNA processing, as shown in the case of TDP-43 autoregulation (Koehler *et al*, 2022). This notion is supported by Hallegger et al., who also showed that TDP-43 phase separation determines the recruitment to RNA transcripts with specific sequence composition (Hallegger *et al*, 2021).

Here, we investigated the role of RNA-mediated TDP-43 self-assembly through oligomerization and phase separation in regulating cellular distribution, and examined the relative contribution of each of these interactions to the native molecular weight of these RNP complexes. We demonstrate that RNA binding and multi-domain interactions form large macromolecular assemblies that act in combination to maintain TDP-43 nuclear localization by preventing nuclear exit. Our work highlights important mechanisms regulating TDP-43 cellular homeostasis by showing that these macromolecular complexes increase nuclear retention in a size-dependent manner and are tightly regulated by RNA-mediated phase separation.

## RESULTS

### TDP-43 multivalent interactions are necessary for TDP-43 nuclear localization

Mounting evidence, including our previous studies, indicate that RNA binding increases nuclear TDP-43 localization (Ayala *et al*., 2008; Duan *et al*., 2022). In addition, our recent work showed that RNA binding, in the presence of multivalent interactions, induces biomolecular condensation or LLPS (Grese *et al*., 2021). We therefore asked whether these RNA-bound complexes play a role in TDP-43 nuclear retention by investigating how disruptions in their assembly alter cellular distribution. We analyzed site-specific mutations in different TDP-43 domains previously established to impair self-assembly (**Fig. 1A**), using an engineered human cell line stably expressing a single copy of hemagglutinin (HA)-tagged wild-type (WT) or TDP-43 variants upon induction with tetracycline (Tet) (Koehler *et al*., 2022). This collection of human embryonic kidney isogenic cell lines (HEK293^HA-TDP-43^) expressed WT and mutant TDP-43 at close to endogenous levels. The small HA tag was specifically used to minimize size-dependent disruption of cellular trafficking. To measure nuclear-cytoplasmic distribution, nucleus and cytoplasm were biochemically separated after 72 h of transgene induction. TDP-43 levels in nuclear and cytoplasmic fractions were measured by immunoblotting (**Fig. 1B, Fig. S1**) and expressed as the nuclear-cytoplasmic ratio (N/C) (**Fig. 1C**). Lamin A/C and tubulin were used to normalize samples and to determine the integrity of nuclear and cytoplasmic fractions for each experiment. The same assays were previously utilized to show that TDP-43 predominantly localizes in the nucleus (Ayala *et al*., 2008). The anti-TDP-43 antibody detected low levels of endogenous TDP-43 upon HA-WT TDP-43 expression (**Fig. 1B**), which is consistent with TDP-43 autoregulatory function (Ayala *et al*, 2011; Polymenidou *et al*., 2011). A mutant disrupting the nuclear localization signal (NLS^mut^, K^82^RK^84^/K^95^VKR^98^ to A^82^AA^84^/A^95^VAA^98^) (Winton *et al*., 2008) was used as a control (**Fig. 1A**). Each independent replicate was performed in parallel with WT-expressing cells, and the N/C was calculated relative to WT (**Fig.1C**). As expected, NLS^mut^ levels increased in the cytoplasm, resulting in a 50% reduction in N/C, relative to WT. The immunoblot also showed loss of autoregulation as seen by the higher levels of endogenous TDP-43 in these cells compared to WT (**Fig. 1B**). Endogenous TDP-43 nuclear-cytoplasmic distribution in NLS^mut^-expressing cells was not affected by the presence of the mutant copy. In agreement with our previous results (Ayala *et al*., 2008; Duan *et al*., 2022), nuclear retention was reduced by mutations in the RNA recognition motifs (RRMs) that greatly impair specific RNA binding affinity F147/149/229/231L (F4L) (Buratti & Baralle, 2001; Grese *et al*., 2021). Next, we probed the role of NTD-driven oligomerization by disrupting interactions previously shown to be important for self-assembly (Afroz *et al*., 2017). The E17R substitution is sufficient to inhibit oligomerization (Wang *et al*, 2018) and impair RNA-driven LLPS (Koehler *et al*., 2022). We found that E17R significantly reduced N/C by approximately 70%, relative to WT. Notably, this effect was even more pronounced than loss of the nuclear localization signal (NLS^mut^). The interaction between Glu17 and Ser48 from two independent NTDs is required for oligomerization (Afroz *et al*., 2017). Ser48 is subject to phosphorylation (Hornbeck *et al*, 2015; Rigbolt *et al*, 2011; Wang *et al*., 2018) and this is predicted to inhibit oligomerization as observed by the decrease in TDP-43 dimerization and LLPS in the presence of the phosphomimetic S48E substitution (Wang *et al*., 2018). Consistent with the effect of E17R, we observed that S48E significantly reduced TDP-43 N/C by approximately 50% compared to WT. In contrast, the phospho-null mutant S48A, used as control, showed a moderate yet not significant increase in N/C relative to WT. Importantly, these results suggest that phosphorylation at Ser48 may regulate TDP-43 nuclear localization. We also found that deletion of the conserved region (ΔCR) in the C-terminal domain decreased the relative N/C by approximately 50%. Altogether, our observations suggest that RNA-dependent multivalent interactions through the NTD and conserved α-helix at the C-terminus strongly promote TDP-43 nuclear localization.

To distinguish whether enhanced cytoplasmic localization of TDP-43 mutants (**Fig. 1C**) was caused by defects in nuclear import, or increased nuclear efflux, we performed heterokaryon assays using WT and mutant HEK293^HA-TDP-43^ cells. HEK293^HA-TDP-43^ cells were fused with NIH3T3 mouse cells resulting in multinucleated cells (**Fig. 2A**). Within one hour of outer membrane fusion, WT HA-TDP-43 was observed in HEK293 nuclei (donor) as well as in mouse nuclei (acceptor) of heterokaryon cells, consistent with our previous findings (Ayala *et al*., 2008) (**Fig. 2B**). Mutants E17R, F4L and ΔCR were also present in acceptor nuclei, indicating that the mutations did not impair nuclear import. Interestingly, we observed that at this time point WT TDP-43 levels in the donor nuclei remained greater compared to the acceptor, based on the fluorescence signal. In contrast, E17R, F4L and ΔCR showed similar levels of HA-TDP-43 fluorescence in donor and acceptor nuclei. To further investigate these differences, we quantified the fluorescence ratio of donor vs. acceptor nuclei (**Fig. 2C**). WT TDP-43 donor to acceptor ratio was approximately 1.5, suggesting that at 1 h post heterokaryon formation, TDP-43 exits donor nuclei and is imported into the acceptor, but that a significant amount of protein, 50% more, is retained in donor nuclei. The mutants, in contrast, showed significant reductions of donor to acceptor ratio (approximately 50-25%), relative to WT. These results indicate that the increased cytoplasmic levels of E17R, F4L and ΔCR, shown in figure 1, are not the result of impaired nuclear import. Rather, disruption of TDP-43 self-assembly through the NTD and CR as well as RNA binding deficiency increase the rate of TDP-43 egress from the nucleus.

**Figure 2.**
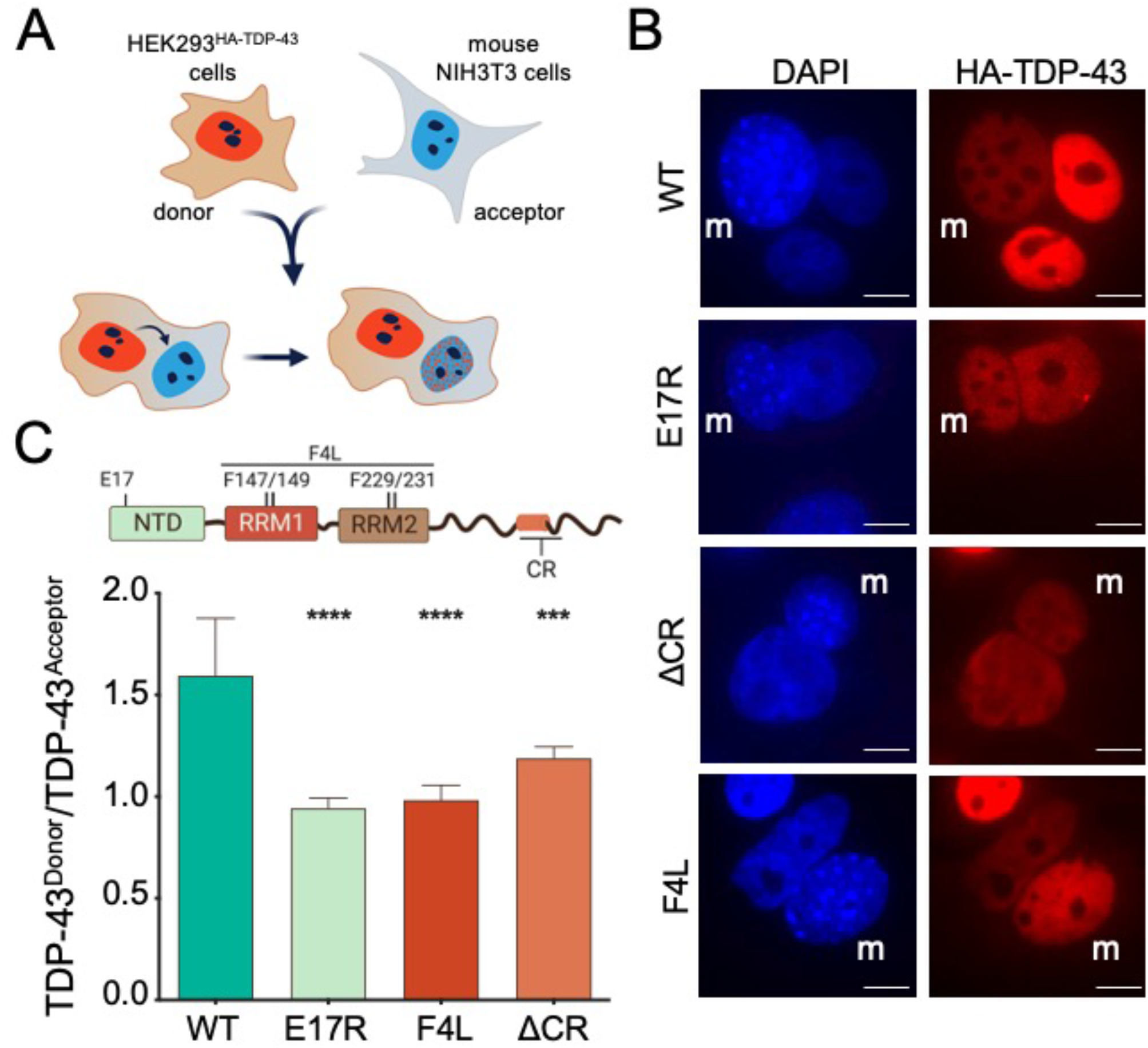
Disruption of RNA binding or TDP-43 self-assembly accelerates TDP-43 nuclear efflux. **A)** Heterokaryon assays in which HEK293^HA-TDP-43^ cells are fused to mouse NIH3T3 cells. HEK293^HA-TDP-43^ nuclei (donor) and HA-TDP-43 signal are depicted in red, mouse NIH3T3 nuclei (acceptor) in blue. **B)** Immunofluorescence of heterokaryons highlighting mouse nuclei (m). Scale, 10 μm. **C)** Quantified HA-TDP-43 signal ratio of HEK293^HA-TDP-43^ (donor) to mouse (acceptor) nuclei of heterokaryons. Mean ± SD shown for 7 independent replicates, 50 or more heterokaryons were quantified, ***p<0.001, ****p<0.0001, one-way ANOVA with Tukey’s test compared to WT.

### RNA binding and multi-domain TDP-43 interactions prevent nuclear exit synergistically

To define the relative contribution of NTD, CR-driven assembly and RNA binding to TDP-43 nuclear retention, we investigated the role of RNA on the localization of NTD and ΔCR mutants. For these experiments we measured nuclear and cytoplasmic TDP-43 levels in HEK293^HA-TDP-43^ cells using confocal high-content imaging (Duan *et al*., 2022). In agreement with the biochemical fractionation assays, E17R, F4L and ΔCR showed significant reductions of TDP-43 N/C, compared to WT (**Fig. 3A**). F4L caused a significantly greater loss of nuclear localization relative to E17R and ΔCR, highlighting a central role of RNA binding in maintaining high nuclear TDP-43 concentration. Treatment with the pan-transcriptional inhibitor Actinomycin D (Act D) reduced WT nuclear localization (**Fig. 3B**), as previously observed (Ayala *et al*., 2008; Duan *et al*., 2022). Similarly, the specific RNA polymerase II inhibitor NVP2 (Olson *et al*, 2018) decreased WT TDP-43 N/C by approximately 40%, indicating that the observed changes are specific to decreased transcription. We found that Act D and NVP2 treatment significantly decreased E17R and ΔCR N/C by approximately 45 and 20%, respectively, compared to untreated control cells. In contrast, localization of F4L was unaffected, consistent with the idea that transcriptional inhibition alters TDP-43 localization by decreasing the abundance of nuclear pre-mRNAs available for TDP-43 to bind. We then transfected cells with increasing amounts of (GU)_6_ RNA oligonucleotides carrying a phosphorothioate backbone and 2’-O-methyl modifications to prevent degradation by RNases (reviewed in (Bennett & Swayze, 2010)). The (CA)_6_ RNA sequence which shows no significant TDP-43 binding (Buratti & Baralle, 2001) was used as non-specific control. Efficient cellular uptake and nuclear import of these RNA molecules was previously quantified (Duan *et al*., 2022). In agreement with Duan et al., (GU)_6_ RNA induced a dose-dependent efflux of WT TDP-43 to the cytoplasm (**Fig. 3C**), while the non-specific RNA did not have an effect. E17R and ΔCR mutant N/C was decreased by (GU)_6_ in the same manner as WT, suggesting that increasing levels of GU-rich RNA further impair nuclear retention of TDP-43 mutants with defects in self-assembly. On the other hand, F4L nuclear-cytoplasmic distribution was insensitive to the presence of (GU)_6_ RNA, as expected. Our results indicate that RNA binding, NTD oligomerization and CR-mediated self-assembly together contribute to TDP-43 nuclear retention and that the sum of these interactions is required to prevent protein efflux to the cytoplasm.

**Figure 3.**
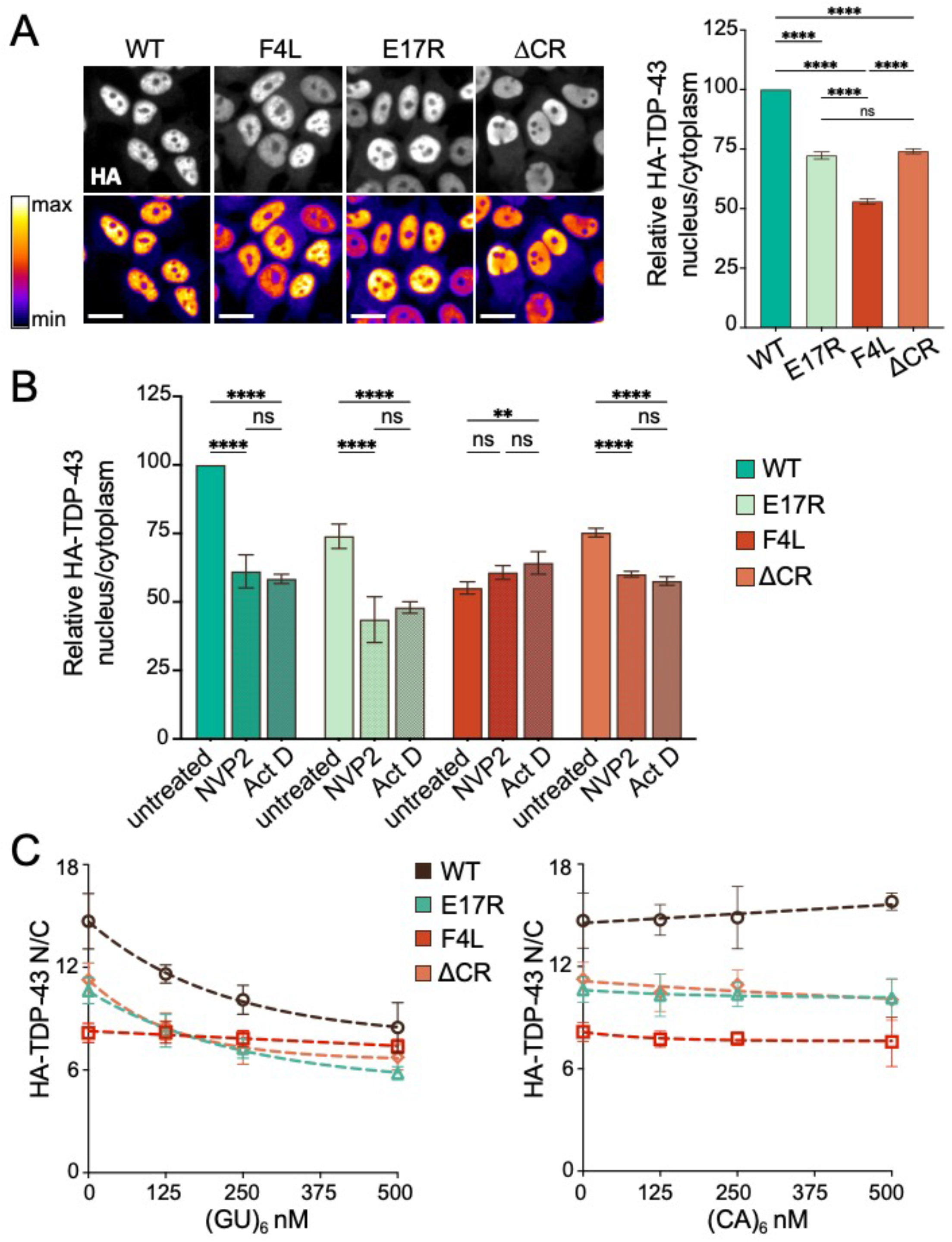
RNA binding and TDP-43 self-assembly exert independent and additive effects on TDP-43 nuclear localization. A) HEK293^HA-TDP-43^ cells expressing HA-tagged WT and mutant TDP-43 probed with HA antibody (top: immunofluorescence, bottom: pseudo-color look-up table). The intensity histogram for each image was independently maximized across the full range. Scale bar, 20 µm. Graph shows automated quantification of the TDP-43 nuclear/cytoplasmic ratio (N/C) at steady state, normalized to WT. Mean ± SD is shown for ∼1600 cells/well in 7 independent replicates. ns, not significant, ****p < 0.0001, one-way ANOVA with Tukey’s test. B) HA-TDP-43 N/C quantified 2 h-post transcriptional blockade with actinomycin D (Act, 1μM) or NVP2 (250 nM) versus DMSO control, normalized to WT control cells. Mean ± SD is shown for ∼1700 cells in 4 independent replicates. ns, not significant, **p < 0.01, ****p < 0.0001, two-way ANOVA with Tukey’s test. C) HA-TDP-43 N/C quantified 5 h post-transfection with increasing concentrations of (GU)_6_ or (CA)_6_ oligonucleotides. Mean ± SD is shown for ∼1700 cells/well in 5 independent replicates.

### Nuclear TDP-43 condensate formation promotes nuclear retention

RNA binding proteins, including TDP-43 and related ALS/FTD-associated proteins, undergo phase separation and condensate formation as purified components and in cells (Mackenzie *et al*, 2017; Molliex *et al*, 2015; Murakami *et al*, 2015; Patel *et al*, 2015). The conserved α-helical region within the C-terminal domain of TDP-43 is essential for promoting phase separation in the context of the isolated domain and full-length protein (Conicella *et al*., 2016; Hallegger *et al*., 2021). We showed that this region is also necessary to form RNA-mediated TDP-43 condensates (Koehler *et al*., 2022). To determine whether the decrease in nuclear localization upon deletion of the CR (**Fig. 2C**) is caused by defects in phase separation, we measured TDP-43 nuclear-cytoplasmic distribution under conditions that inhibit condensate formation. 1,6-hexanediol (1,6-HD) is widely used to inhibit phase separation (Kroschwald, 2017) and impairs droplet formation of purified TDP-43 (Mann *et al*., 2019; Schmidt *et al*, 2019). 1,6-HD also decreases the formation of nuclear TDP-43 foci in cells (Hallegger *et al*., 2021), suggesting that it reduces TDP-43 RNP condensates. We treated cells with 5% 1,6-HD for 5 minutes and observed a 5-fold decrease in N/C of endogenous TDP-43, compared to non-treated control cells (**Fig. 4A**). We considered that 1,6-HD may also disrupt the phenylalanine-glycine (FG) permeability barrier in the central channel of the nuclear pore complex (Ribbeck & Gorlich, 2002). This effect could also contribute to the increased nuclear efflux of TDP-43 upon 1,6-HD treatment. To test disruption of TDP-43 condensate formation more specifically, we introduced mutations in the CR to modulate phase separation by altering α-helix structural properties (**Fig. 4B**) (Conicella *et al*., 2016; Hallegger *et al*., 2021). A326P, M337P and the ALS-linked mutation M337V are helix breaking mutants that result in aberrant CTD self-assembly and impaired LLPS (Conicella *et al*., 2016; Grese *et al*., 2021; Koehler *et al*., 2022). Moreover, these single-site substitutions closely resemble the inhibition properties of ΔCR on the formation of TDP-43-rich nuclear foci and phase separation-dependent RNA processing of TDP-43 targets in cells (Hallegger *et al*., 2021). We found significant decrease in nuclear TDP-43 localization with each of these mutations (**Fig. 4B**), consistent with the effect of 1,6 HD treatment and deletion of the entire CR in figure 1C. Altogether, these data strongly suggest that nuclear condensate formation plays a key role in regulating nuclear-cytoplasmic TDP-43 trafficking by promoting nuclear localization.

**Figure 4.**
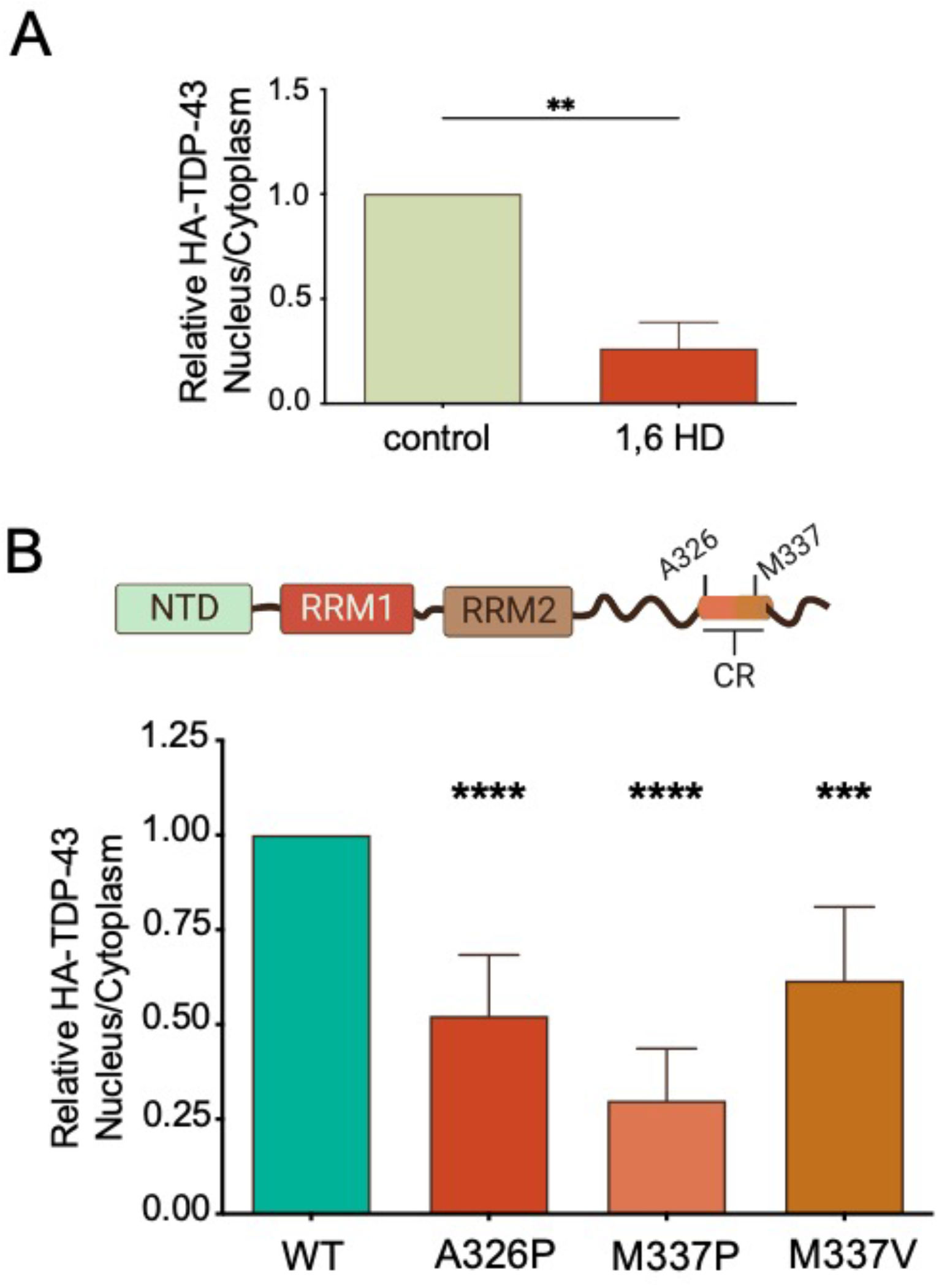
Inhibition of phase separation disrupts TDP-43 nuclear retention. A) Relative nuclear to cytoplasmic signal ratio (N/C) of endogenous TDP-43 in HEK293 cells treated with 1,6 hexanediol (HD, 5% for 5 min) quantified from immunoblotting (Fig. S2A), expressed relative to non-treated control. Mean ± SD is shown for 6 independent replicates. B) HEK293^HA-TDP-43^ cells expressing HA-tagged WT and A326P, M337P and M337V mutants located in the conserved α-helical region (CR). Quantification of N/C of HA-TDP-43 relative to WT in each replicate from immunoblotting (Fig. S2B). Mean ± SD is shown for 6 independent replicates. ***p<0.001, ****p < 0.0001, one-way ANOVA with Tukey’s test.

### The size of RNA-bound TDP-43 macromolecular complexes controls nuclear retention

Previous evidence indicates that TDP-43 nuclear efflux is largely a passive, size dependent process, whereby larger fusion constructs reduce TDP-43 export (Ederle *et al*., 2018; Pinarbasi *et al*., 2018). TDP-43 forms complexes of a wide range of sizes in cells, from monomers to large higher-order complexes greater than 1.5 MDa, as measured by size exclusion chromatography (SEC) (Guo *et al*, 2011; Mallam *et al*, 2019). Based on these findings, we tested whether the contribution of TDP-43 RNA-mediated complexes to nuclear retention is size-dependent, requiring the formation of large macromolecular complexes. We analyzed the size distribution of TDP-43 in HEK293^HA-TDP-43^ cell lysates expressing WT and mutant TDP-43 using gel filtration chromatography. Of note, the use of these isogenic cell lines expressing TDP-43 variants at endogenous-like levels was designed to avoid unintended effects of TDP-43 overexpression. The relative HA-TDP-43 levels in each eluted fraction was analyzed by immunoblotting to compare the approximate size distribution of WT vs mutant TDP-43 assemblies (**Fig. 5A, Fig S3**). We observed a broad profile of WT TDP-43 distribution starting at the estimated 45 kDa fraction to 700 kDa, which is the upper detection limit of the SEC column. The highest level of detected WT protein eluted at approximately 115 kDa and this was set as 100% (**Fig. 5B**). Disruption of NTD-mediated oligomerization, E17R, showed a shift to higher elution volume compared to WT, whereby the peak of highest concentration moved to smaller estimated molecular weight, between 45 and 72 kDa (**Fig. 5C**). In addition, TDP-43 levels in the fractions corresponding to higher molecular weight complexes were markedly decreased compared to WT. These results are consistent with previous cross-linking assays indicating a loss of multimeric TDP-43 species in cells upon disruption of NTD-mediated oligomerization (Afroz *et al*., 2017). Similar behavior was observed with the RNA-binding-deficient F4L mutant. The most abundant F4L fractions shifted to smaller estimated sizes compared to WT together with a dramatic sharpening of the elution profile. An even greater shift in size distribution was observed with ΔCR, showing an elution peak of estimated 45 kDa. In stark contrast with WT, little or no detection of ΔCR was observed above the estimated 115 kDa fraction. Based on the estimated size separation in our experiments, the altered behavior of ΔCR is unlikely to be caused by the 20 amino acid difference between this construct, WT and the other mutants analyzed here. To further test the role of CR-mediated assembly and LLPS on the size of TDP-43 complexes, we studied the effect of A326P. In agreement with ΔCR, A326P also reduced the size of the most abundant TDP-43 complexes, eluting at the estimated 45 kDa fraction. These results are consistent with an important role of CR-mediated homotypic assembly and LLPS in forming higher-order RNP bodies. Collectively, our SEC experiments provide biochemical evidence that disruption of NTD oligomerization, RNA binding and phase separation decrease the size of TDP-43-rich complexes by impairing formation of the largest macromolecular species in cells. In addition, our data strongly suggest that the decreased ability to form these higher order macromolecular assemblies reduces TDP-43 nuclear retention by increasing the population of protein freely diffusing to the cytoplasm.

**Figure 5.**
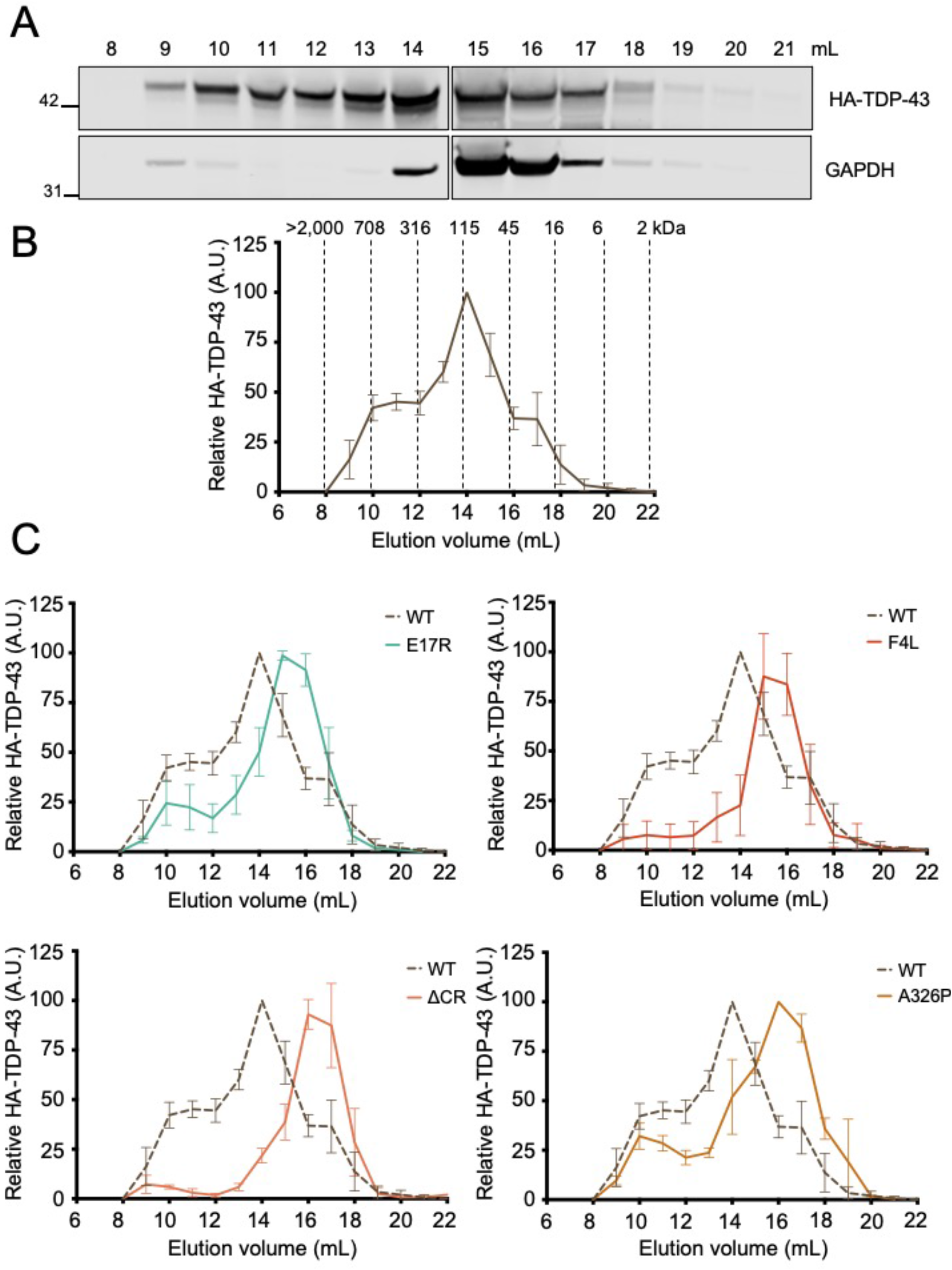
Disruption of RNA binding, NTD oligomerization and phase separation inhibits the assembly of large macromolecular TDP-43 complexes in cells. A) Immunoblot detecting HA-TDP-43 levels in fractions eluted from gel filtration size exclusion chromatography of HEK293^HA-TDP-43^ cell lysate. Membranes were probed with HA antibody and GAPDH as control. B) WT HA-TDP-43 levels quantified in each elution fraction expressed as percent relative to the most abundant TDP-43 fraction. Mean ± SD is shown for 4 independent replicates. The estimated molecular weight is shown for each fraction based on calibration experiments. C) Comparison of elution profiles for WT (dashed) vs. mutant (solid) TDP-43. Mean ± SD is shown for ≥3 independent replicates.

## DISCUSSION

The normal distribution of TDP-43 between nuclear and cytoplasmic compartments, in which the protein is predominantly found in the nucleus, plays a central role in physiological function. Disruption of homeostasis is associated with pathogenesis in neurodegenerative disorders, wherein pathological decrease in nuclear TDP-43 and elevated protein levels in the cytoplasm are linked to TDP-43 loss of function and aggregation, respectively (Arai *et al*., 2006; Liu *et al*., 2019; Neumann *et al*., 2006). TDP-43 is recruited to RNA and protein-rich ribonucleoprotein (RNP) complexes through RNA binding, TDP-43 self-assembly and condensate formation/phase separation (Bentmann *et al*, 2012). The mechanisms by which these complexes regulate TDP-43 function and homeostasis are only beginning to be explored. Recently, we showed that RNA-mediated TDP-43 phase separation depends on multivalent interactions involving N-terminal domain (NTD) oligomerization and assembly through an α-helical conserved region at the C-terminus (CR) (**Fig 1A**) (Grese *et al*., 2021; Koehler *et al*., 2022), which is essential for TDP-43 condensation in vitro and in cells (Conicella *et al*., 2016; Hallegger *et al*., 2021; Schmidt & Rohatgi, 2020). Here, we find that disruption in any of the assembly points necessary to form RNA-mediated TDP-43 complexes is sufficient to markedly increase cytoplasmic localization. Thus, TDP-43 recruitment into RNP bodies plays a critical function in maintaining nuclear retention.

Based on emerging evidence, RNA binding plays a key role in TDP-43 molecular dynamics and proteostasis. We recently showed that specific RNA binding modulates TDP-43 phase separation properties (Grese *et al*., 2021) and that formation of these complexes is essential in regulating TDP-43 target expression, such as its own transcript during autoregulation (Hallegger *et al*., 2021; Koehler *et al*., 2022). The importance of RNA in TDP-43 nuclear localization was previously demonstrated with RNA binding-deficient mutants and by showing that inhibition of RNA transcription promotes cytoplasmic localization (Ayala *et al*., 2008; Duan *et al*., 2022). In addition, Duan et al. recently quantified the effect of TDP-43-RNA interactions in cells and showed that inhibition of splicing increases TDP-43 nuclear levels probably by increasing the retention of introns with TDP-43 recruitment sites (Duan *et al*., 2022). In addition, small GU-rich RNA oligonucleotides, which bind TDP-43 with nanomolar affinity (Ayala *et al*., 2005), promote the cytoplasmic egress of TDP-43 (Duan *et al*., 2022). These data suggest that these RNA molecules promote the release of nuclear TDP-43 by competing with transcripts for TDP-43 binding. Here, we find that mutants deficient in NTD and CR-driven interactions are similarly affected by (GU)_6_ RNA compared to WT (**Fig. 3C**). Thus, our observations indicate that RNA binding, NTD and CR interactions contribute synergistically in maintaining nuclear localization. Importantly, we and others showed that specific RNA interactions prevent TDP-43 aggregation (French *et al*, 2019; Mann *et al*., 2019) and promote liquid-like behavior of TDP-43 condensates (Grese *et al*., 2021). These results collectively suggest that RNA binding plays multiple essential roles in maintaining TDP-43 function. Specific RNA binding increases nuclear TDP-43 retention by mediating the formation of RNP assemblies. In addition, RNA interactions increase solubility in the nucleus despite the higher protein concentration in this compartment. This idea is consistent with previous observations that solubility and liquid properties of the related protein FUS are maintained by the abundance of RNA in the nucleus (Maharana *et al*., 2018).

Our data agree with recent work from the Polymenidou lab published while our manuscript was in preparation (Perez-Berlanga *et al*, 2023). Using cell-based models, including human neurons, Pérez-Berlanga et al. show that NTD-mediated oligomerization and RNA binding act cooperatively to increase nuclear localization and that both interactions are important to prevent TDP-43 aggregation. Here, we extend beyond these findings by showing that condensate formation is required for the assembly of complexes that increase TDP-43 nuclear retention (**Fig 4**), highlighting the importance of multivalent interactions in TDP-43 function. In addition, our heterokaryon assays demonstrate that the defects in nuclear localization observed upon disruption of TDP-43 RNA binding, oligomerization or phase separation are not caused by diminished nuclear import (**Fig 2**). Instead, our results suggest that disruption of these interactions accelerate protein efflux to the cytoplasm. Moreover, our size exclusion chromatography studies are the first to biochemically correlate the ability of TDP-43 to form RNA-bound condensates with the formation of large higher-order macromolecular complexes in cells (**Fig 5**). Based on our observations, we propose that TDP-43 nuclear retention is largely dependent on the formation of large RNP granules, reducing the population of smaller species available for passive export to the cytoplasm (**Fig 6**). These results are important because they support a size-dependent nuclear retention mechanism of RNA-bound TDP-43 condensates and illustrate multiple possible mechanisms to modulate TDP-43 nucleocytoplasmic distribution. Furthermore, our observations raise the intriguing possibility that the cellular dynamics of other related RNA-associated proteins may be controlled in similar fashion.

**Figure 6.**
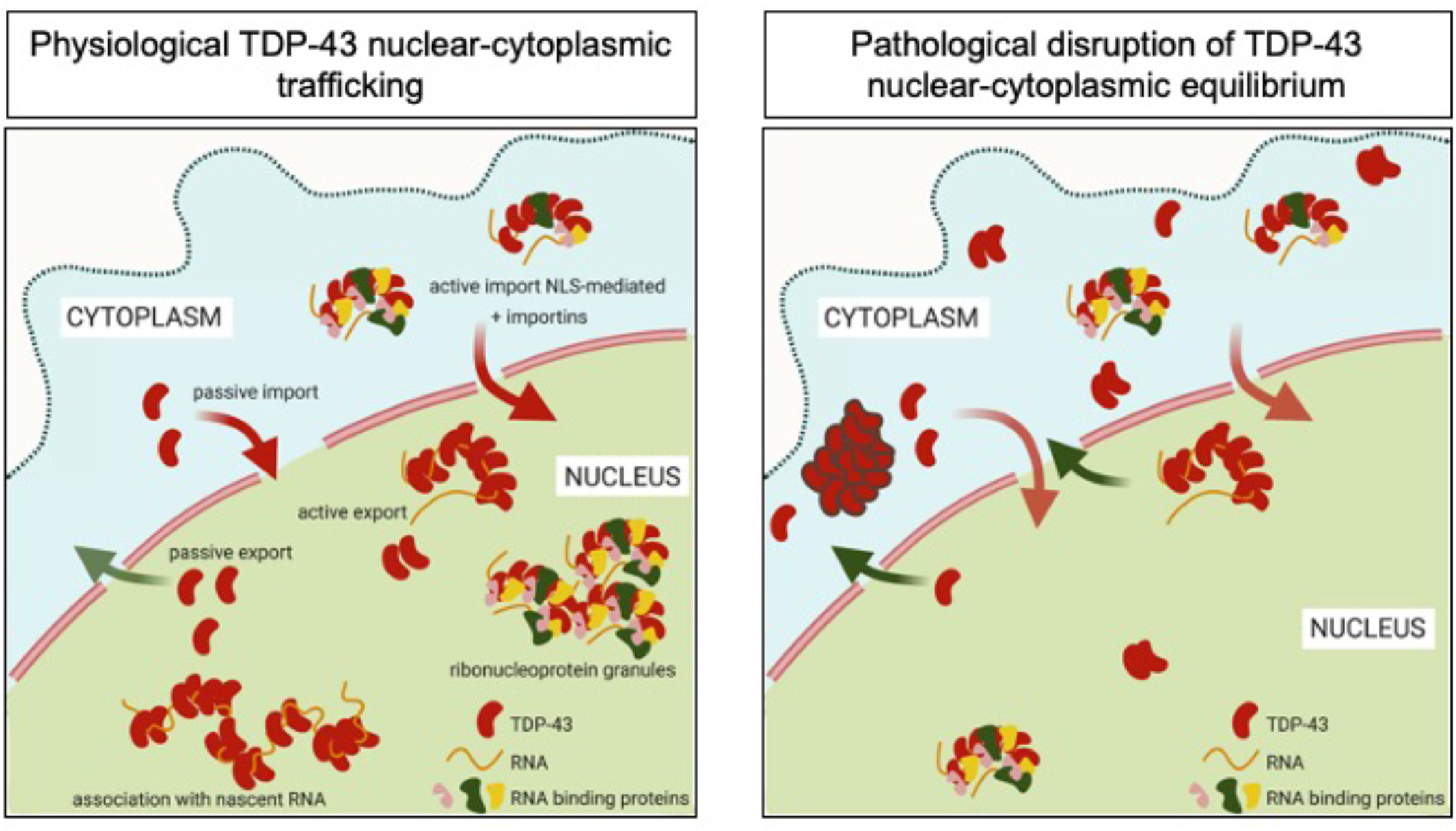
TDP-43 nuclear localization depends on the assembly of RNP complexes mediated via multivalent interactions. Ribonucleoprotein TDP-43 complexes in the nucleus contribute to TDP-43 nuclear localization through the formation of higher-order macromolecular assemblies unable to passively diffuse to the cytoplasm. Under aberrant conditions, disruption of RNA binding or self-assembly increases levels of smaller TDP-43 species inducing greater cytoplasmic localization. Chronic defects in RNP assembly may increase TDP-43 misfolding and aggregation in the cytoplasm.

Condensate formation plays a central role in RNP granule assembly and function (Shin & Brangwynne, 2017) and our work suggests that it promotes TDP-43 nuclear localization, as seen by deleting and/or disrupting the CR in the C-terminus. This region also interacts with different RNA binding proteins including hnRNPs A1, A2, A3, B1 and C (D’Ambrogio *et al*, 2009). Therefore, association with these hnRNPs may also contribute to RNP granule formation and nuclear retention, and this possibility requires further investigation. We also observe a significant decrease in nuclear localization in the presence of A326P, M337P and the ALS-linked mutation M337V and a reduction in the size of TDP-43 complexes with the A326P mutation, similar to the effect of ΔCR. We predict that these single-site substitutions do not abolish TDP-43 association with the partner hnRNPs. On the other hand, the substitutions significantly disrupt TDP-43 phase separation, as seen in vitro and in cells (Conicella *et al*., 2016; Hallegger *et al*., 2021). Therefore, we posit that CR contributes to TDP-43 nuclear localization primarily through homotypic assembly and phase separation.

Collectively, our findings suggest the intriguing possibility that functional defects previously observed with oligomerization-deficient and other mutants/conditions affecting TDP-43 self-assembly may be at least partially caused by decreased nuclear localization. Since the structural characterization of TDP-43 NTD-mediated oligomerization (Afroz *et al*., 2017; Wang *et al*., 2018), this interaction has been increasingly shown to play a central role in cellular functions. NTD-driven oligomerization is required for TDP-43 function in alternative splicing regulation (Afroz *et al*., 2017; Wang *et al*., 2018), phase separation (Grese *et al*., 2021; Wang *et al*., 2018), autoregulation (Koehler *et al*., 2022) and in preventing R-loop accumulation (Wood *et al*., 2020). Here, we find that impaired oligomerization greatly reduces nuclear TDP-43 localization (**Fig 1C**), indicating that loss of function due to inhibition of NTD-mediated oligomerization may be caused by multiple mechanisms. These defects may include decreased self-assembly upon RNA target engagement, reduced RNP body formation, and decreased nuclear concentration. Importantly, our work suggests that TDP-43 nuclear efflux to the cytoplasm may be regulated by posttranslational modifications, mainly through phosphorylation at Ser48 in the NTD. We show that the phosphomimetic mutation S48E, which inhibits NTD-driven oligomerization (Wang *et al*., 2018), decreases nuclear TDP-43 localization by almost 50%. Thus, phosphorylation at Ser48 may normally tune TDP-43 nuclear-cytoplasmic distribution and aberrant activation of the phosphorylation pathways, which remain unknown, may contribute to increased cytoplasmic TDP-43 accumulation.

Together, our studies highlight multiple mechanisms that control TDP-43 RNP assembly and nuclear-cytoplasmic distribution through different types of interactions mediated by distinct TDP-43 domains, including binding to various RNA sequences with different affinities. This idea also implies that dysfunction of these regulatory factors may result in abnormal or pathological cytoplasmic TDP-43 accumulation. Therefore, these mechanisms may be targeted for therapeutic intervention to restore TDP-43 homeostasis and function.

## Acknowledgements

This work was supported in part by National Institutes of Health (NIH) National Institute for Neurological Disorders and Stroke (NINDS) and National Institute on Aging (NIA) grants R01 NS114289 (Y.M.A.) and R01NS123538 (L.H.); and the Department of Defense CDMRP/ALSRP W81XWH-20-1-0241 (Y.M.A.). We thank Lydia Koehler for her technical help in creating stable cell lines and performing the initial experiments in these studies.

We thank Dr. Joel Eissenberg and Dr. Alessandro Vindigni for their careful editing of the manuscript.

## MATERIALS AND METHODS

Chemicals were purchased from Sigma Aldrich unless otherwise specified.

### Cloning, cell culture and stable cell production

Generation of HEK293^HA-TDP-43^ stable cell lines, expressing HA-tagged WT, E17R, S48A, S48E, M337V and ΔCR constructs and stable cell lines was previously described (Koehler *et al*., 2022). Mutants A326P and M337P were generated by site-directed mutagenesis as previously described(Grese *et al*., 2021) using the primers listed in Supplementary Table 1. Briefly, the HEK293-derived cells Flp-In^TM^ T-Rex^TM^ 293 Cell Line (ThermoFisher Scientific) were stably transfected to express HA-TDP-43 upon induction with tetracycline (1 μg/mL). Cells were grown and maintained in DMEM (Dulbecco’s Modified Eagle’s Medium – High Glucose, Corning) supplemented with 10% FBS (Fetal Bovine Serum) and incubated in a humid atmosphere at 37 °C and 5% CO_2_. Expression of HA-tagged TDP-43 construct was induced at 30% confluence.

### Cell fractionation

Stable cell lines cultured in 10 mm dishes (TPP) were induced with 1 μg/mL tetracycline for 72 h prior to harvesting and were pelleted through centrifugation at 300*g* for 5 minutes. Cell pellets were washed twice with PBS (phosphate buffered saline) and gently resuspended in 2 mL of Lysis Buffer A (10 mM HEPES pH 7.9, 1.5 mM MgCl_2_, 10 mM KCl, 1 mM TCEP, protease inhibitor cocktail, 10 mM NaF, 1.5 mM orthovanadate, 10 mM β-glycerol phosphate). Resuspended samples were incubated for 5 min on ice and Dounce homogenized with 10 strokes (Pyrex, No 7727). Samples were pelleted at 200*g* for 5 min and supernatants were collected as cytoplasmatic fractions. The pellets were rinsed in 2 mL of PBS and pelleted at 200*g* for 5 min. The pellets were resuspended in 500 μL of PBS to obtain the nuclear fraction. All fractionation steps were carried out at at 4°C using cold buffers. Fractions were mixed with Laemmli Buffer (0.2 M Tris pH 6.8, 20% Glycerol, 10% SDS, 1.43 M β-mercaptoethanol, 0.05% bromophenol blue) and heated for 5 min at 95°C. Equal volumes were resolved in standard 12% SDS-PAGE and immunoblotted against specified antibodies.

### Heterokaryon assay

HEK293^HA-TDP-43^ cells were grown on glass coverslips and heterokaryon assays were performed as previously described (Ayala *et al*., 2008).

### 1,6-Hexanediol Treatment

5% 1,6-hexanendiol was added to HEK293^HA-TDP-43^ cells cultured at 80 to 90% confluency for 5 minutes prior to harvesting.

### Size exclusion chromatography

HEK293^HA-TDP-43^ cells were harvested at 90% - 100 % confluence following 72 h of tetracycline induction. Cells were washed with PBS and resuspended in lysis buffer (25 mM Tris HCl pH 7.5, 0.1 mM Na_3_VO_4,_ 10 mM MgCl_2_, protease inhibitor cocktail, 150 mM NaCl, 0.5% NP-40, 2 mM DTT, 5 mM β glycerophosphate) and processed using a Dounce homogenizer with 20 strokes (Pyrex, No 7727). Following centrifugation, soluble proteins were applied to the size exclusion gel filtration column (Superdex 200 Increase 10/300 GL, GE Healthcare), using an Akta P-920 Pump FPLC system (Amersham) and Running Buffer: 25 mM Tris-HCl (pH 7.5), 150 mM NaCl. Fractions were collected at 0.5 ml min^−1^. The resulting dilution from input lysate was approximately 60-fold. Fractions were analyzed by standard SDS-PAGE followed by immunoblotting. The approximate molecular weight of each fraction was determined based on the elution volumes of thyoglobulin (670 KDa), γ globulin (158 KDa), ovalbumin (44 KDa), myoglobin (17 KDa), vitamin B12 (13.5 KDa), according to manufacturer instructions.

### High content analysis of RNA-dependent TDP-43 localization

HEK293^HA-TDP-43^ cell lines (WT, F4L, E17R, and delta-CR) were plated on Ultimatrix-coated (1:100, R&D), 96-well optical plastic plates (Ibidi) to achieve ∼75% confluence. Doxycycline (1 μg/mL) was added to culture media at the time of plating, targeting 24 h of induction at the time of fixation. Transcriptional blockade was by NVP2 (250 nM, Tocris) or actinomycin D (1 µM) for 2h. RNA oligonucleotides (IDT, see table) were transfected using Lipofectamine RNAiMax (Thermo Fisher) according to the manufacturer’s instructions except that the lipofectamine amount was doubled based on initial optimization studies (not shown). Cells were fixed for analysis 5 h post-transfection. To determine the localization of HA-tagged TDP-43 and mutants, cells were fixed with 4% paraformaldehyde in PBS, permeabilized and blocked in 0.1% Tx-100 and 10% normal goat serum (Vector Labs), and immunostained with rabbit anti-HA (1:1500, Cell Signaling Technologies) overnight at 4°C in 10% NGS/PBS. After two PBS rinses, Alexa Fluor-labeled secondary antibodies were added for 1 h at room temperature (1:1000, Thermo). Cells were rinsed with PBS containing Hoechst 33342 and transferred to 50% glycerol/PBS for automated imaging and analysis using an ImageXpress Micro Confocal high content microscope with MetaXpress software (Molecular Devices) as described (Duan et al., 2022). A minimum of four independent biological replicates was analyzed per experiment.

## Supplementary Material

**Supplementary figure 1.**
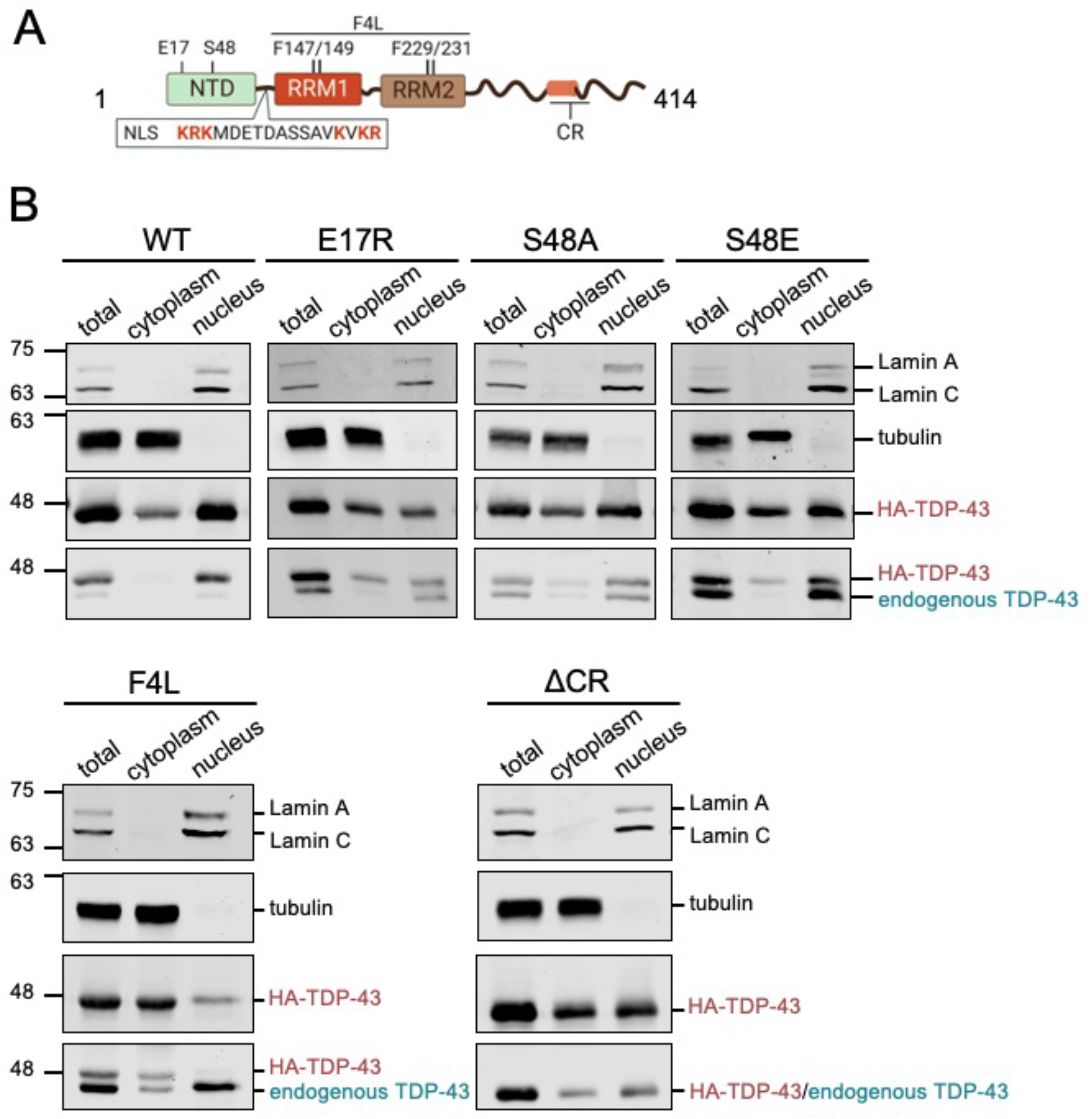
Defects in TDP-43 self-assembly and RNA binding decrease nuclear localization. A) TDP-43 domain organization highlighting targeted mutation sites to inhibit N-terminal domain (NTD) oligomerization, RNA binding and C-terminal assembly through deletion of the conserved α-helical region (CR) in the C-tail (a.a. 316-343). The basic residues (red) in the nuclear localization signal (NLS) were substituted to generate NLS^mut^, used as control. B) TDP-43 levels in nuclear and cytoplasmic compartments were compared after biochemical fractionation. The immunoblots detect HA-TDP-43 and endogenous TDP-43 in total, cytoplasmic and nuclear fractions of the stable HEK293^HA-TDP-43^ cells. Lamin A/C and tubulin were used as nuclear and cytoplasmic controls, respectively.

**Supplementary figure 2.**
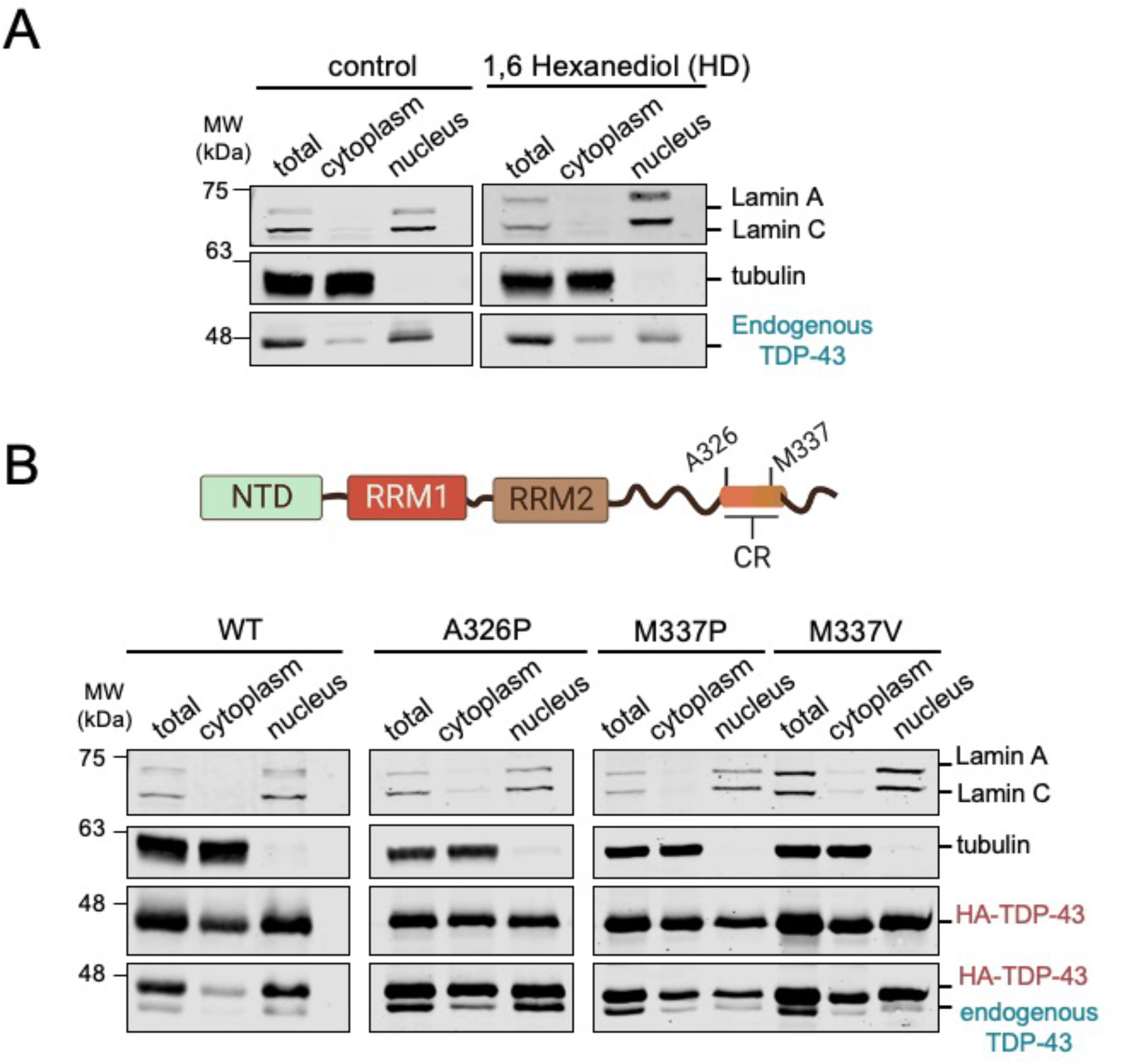
Inhibition of phase separation disrupts TDP-43 nuclear retention. A) Biochemical fractionation of HEK293 cells treated with 1,6 hexanediol (HD, 5% for 5 min), or non-treated cells into nuclear and cytoplasmic compartments. Immunoblots probed for endogenous TDP-43 in total, nuclear and cytoplasmic fractions. Lamin and tubulin are nuclear and cytoplasm controls, respectively. B) Immunoblots of total, nuclear and cytoplasmic fractions from HEK293^HA-TDP-43^ cells expressing phase separation-defective mutants, including ALS-linked M337V.

**Table 1.**
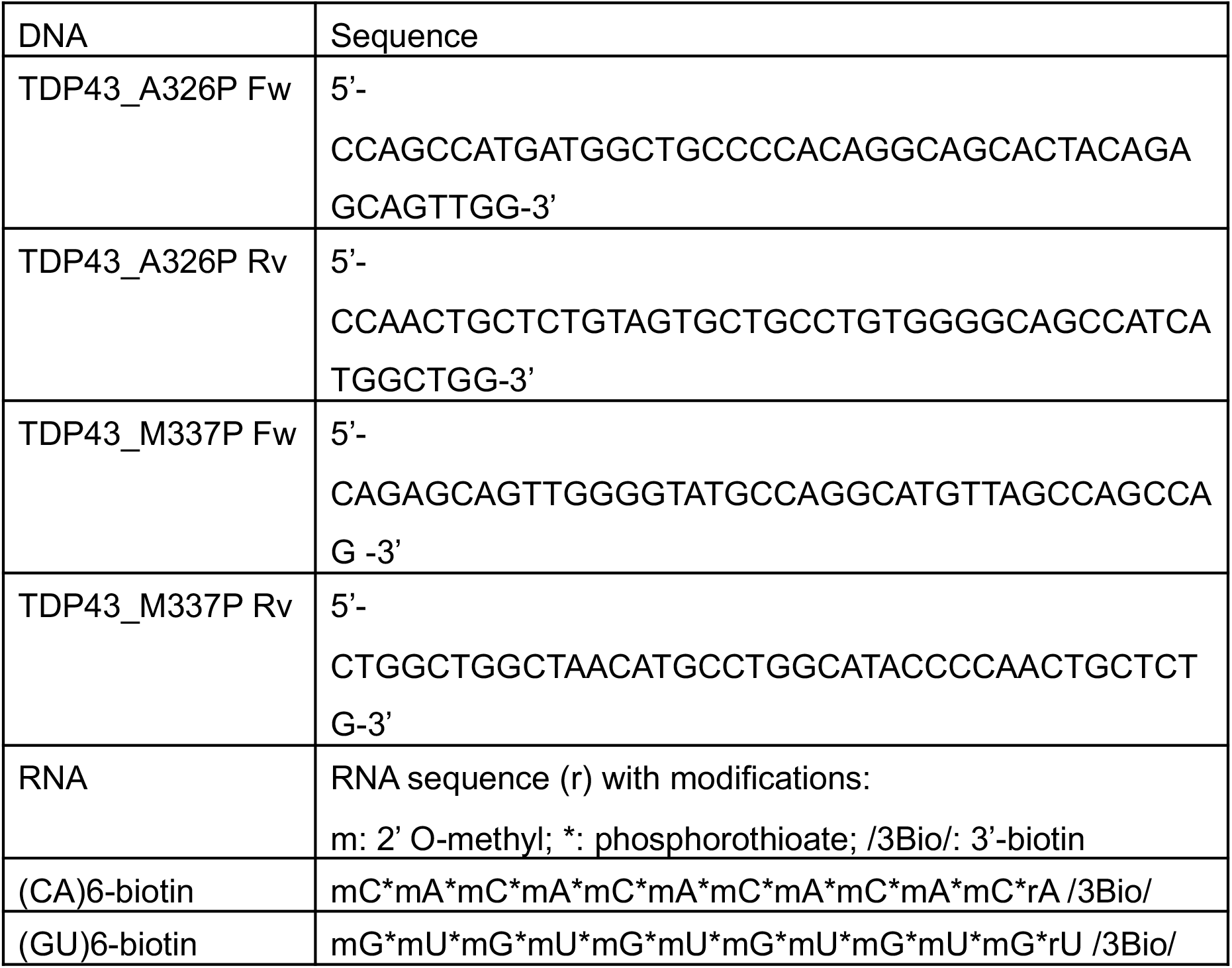
Oligonucleotides used in these studies.

